# COMPARATIVE EFFECTS OF FLAXSEED SUPPLEMENTATION ON HEMATOLOGICAL PARAMETERS, LIPID PROFILE AND IMMUNITY OF MALE RABBIT

**DOI:** 10.64898/2026.04.06.716729

**Authors:** Ayesha Kanwal, Razia Iqbal, Farah Saleem, Amna Kanwal

## Abstract

Flaxseeds have high nutritive value due to the presence of proteins, lignins (SDG), fatty acids, vitamins, dietary fibers, minerals and carbohydrates. This research was conducted to evaluate the effect of distinct doses of flaxseeds on hematological parameters, immunity and lipid profile of male rabbit. In this research, 60 male rabbits were isolated into four groups, three treatment groups T_1_, T_2_ and T_3_ and a control group T_0_, with 15 rabbits in each group. The treatment groups were given 4%, 6% and 8% of flaxseeds per daily diet for 45 days. On 15^th^, 30^th^ and 45^th^ day of experiment, blood samples were collected to examine their hematological parameters. Serum was separated from the collected blood sample to perform ELISA and serum lipid profile test to assess antibody titer and lipid profile of the rabbits respectively. The results indicated a significant reduction in TC, TG, and LDL-C levels in contrast to HDL-C which increased significantly in all treatment groups. Whereas, the statistical analysis of hematological parameters showed an HSD (p≤0.05) in flaxseed treated groups. A maximum level of Hb, WBCs, RBCs, MCHC, MCV, HCT, MCH and differential leukocytes count was recorded in high dose group T_3_ (8% flaxseeds) followed by medium dose group T_2_ (6% flaxseeds) and low dose group T_1_ (4% flaxseeds) respectively. There was a significant rise in antibody titer (p≤0.05) against RHDV (Rabbit Hemorrhagic Disease Virus) comparable to non-treated group. The outcomes illustrated that flaxseeds as nutritional supplement are undoubtedly beneficial to health and prevent various diseases.

**Study contribution:** This research specifically explores how dietary supplementation with flaxseeds, a widely recognized source of omega-3 fatty acids, fiber, and antioxidants, can influence metabolic health and immune function. These findings have significant implications for nutritional interventions aimed at improving cardiovascular health, immune support, and overall well-being, making it highly relevant to the journal’s readership.

The aim of this study was to investigates the dose-dependent effect of flaxseeds on hematological parameters, lipid profile and immunity of male rabbits. Using a controlled experimental design, male rabbits were fed a diet supplemented with varying doses of flaxseeds over a period of 45 days. Key parameters such as total cholesterol, triglycerides, LDL-C, HDL-C, antibody titer, red, white blood cell, platelet counts, Hb, HCT, MCV. MCHC, MCH and differential leukocytes levels were measured to assess the impact of flaxseeds. The results demonstrated that flaxseed supplementation significantly restored lipid profiles by reducing total cholesterol and triglycerides, LDL-C and increasing HDL-C while also enhancing immune function by rising antibody titer and maintaining healthy blood profiles in the subjects.

## Introduction

The nutritive value of the flaxseeds is due to the presence of bioactive compounds like lipids (Alpha linolenic acid - ALA), proteins, vitamins, peptides (cyclic peptides), minerals, lignins, carbohydrates, and mucilage [10, 22]. Apart from their nutritive value, they exhibit anti-inflammatory and immunoregulatory role due to the presence of excessive amount of ALAs or n-3 PUFAs [11]. Flaxseeds also proclaim hypolipidemic effects due to the existence of lignins, that are the prominent phytoestrogens within flaxseeds [20]. One of the abundant lignins (or phytoestrogen) within flaxseed is SDG (Secoisolariciresinol Diglucoside) [14].

As flaxseeds have significant bioactive compounds with the properties that can influence various parameters of lipid profile, blood profile and immunity, it makes good sense to postulate the effect of flaxseeds on these parameters. So, the main purpose of this research work is to investigate its effect of its different doses on lipid profile, hematological parameters and immunity of male rabbit.

### Research objectives

a) To investigate the effect of different doses of flaxseeds on the lipid profile of rabbits. b) To access the effect of various doses of flaxseeds on the hematological parameters and immunity of rabbits

## Materials and method

### Ethical Statement

All procedures and tests applied on rabbits were approved by the ethical committee of the University of Gujrat, Punjab, Pakistan.

All procedures involving animals were conducted in accordance with institutional ethical standards and the ARRIVE (Animal Research: Reporting of In Vivo Experiments) guidelines.

### Research specimen and dose preparation

60 male rabbits (*Oryctolagus cuniculus*) were purchased from Veterinary Research Institute, Ghazi Road Lahore, Punjab, Pakistan during the month of February and exploited in the current experiment. Rabbits weighing 1-1.5 kg were acclimatized before experimentation for 15 days with optimum light (12 hrs light and dark cycle) and 23 ± 2^°^C temperature The ad libitum feed and water was offered occasionally to the rabbits during the whole experiment [7]. 60 rabbits were classified into 4 groups T_0_, T_1_, T_2_, T_3_, 15 rabbits in each group.

Flaxseeds used in the experiment were obtained from the botanical garden of University of Gujrat (UOG) and grinded into fine powder. After grinding, measured quantity of flaxseeds were used to prepare low 4%, medium 6% and high 8% flaxseed’s dose per daily diet of the rabbit. All groups were given different doses of flaxseeds T_0_ (0% flaxseeds-control group), T_1_ (4%), T_2_ (6%) and T_3_ (8%) per daily diet. The flaxseed dose mixed with soaked bread was given orally once a day for 45 days consecutively [28].

All procedures and tests applied on rabbits were approved by the ethical committee of the University of Gujrat, Punjab, Pakistan.

### Blood collection

At 15^th^, 30^th^ and 45^th^ day, 5 rabbits were selected randomly from each group for blood collection to perform CBC, lipid profile test and ELISA. The blood sample was taken from marginal ear vein with aid of sterile syringe following the standard instructions into 3 ml K_3_ EDTA tubes for CBC and gel & clot activator tubes (3 ml) for serum lipid profile test and ELISA [19].

### Complete blood count for hematology

At 15^th^, 30^th^ and 45^th^ day, 2 ml blood samples were collected from selected 5 rabbits of each group in labelled 3 ml K_3_ EDTA tubes for the measurement of hematological parameters hemoglobin (Hb), hematocrit (HCT), red blood cells (RBCs), white blood cells (WBCs), platelets, mean corpuscular volume (MCV), mean corpuscular hemoglobin (MCH), mean corpuscular hemoglobin concentration (MCHC) and differential leukocytes using auto hematology analyzer [27]. Graphical and numerical values were shown on the screen and changes in the results were compared to the control group, everytime at 15^th^, 30^th^ and 45^th^ day.

### Serum extraction

For serum extraction, collected blood in gel tubes (without coagulant) was centrifuged at 1600xg speed at 4^°^C for 15 minutes and later stored at −70^°^C in serum capsules in refrigerator [25].

### Serum lipid profile test for blood chemistry

At 15^th^, 30^th^ and 45^th^ day, from selected rabbits, labelled gel tubes containing serum were taken to perform lipid profile test using Hitachi 704 chemistry analyzer. The lipid profile test provided the levels of serum total cholesterol (TC), triglycerides (TG), high-density lipoprotein (HDL-C), and low-density lipoprotein (LDL-C) of rabbits using commercial kits [15].

### ELISA for humoral immunity

From serum of 5 selected rabbits (of each group), antibody titer was calculated against RHDV (Rabbit Hemorrhagic Disease Virus) at 15^th^, 30^th^ and 45^th^ day via ELISA. By measuring antibody titer, the change in the humoral immunity of the rabbits belonging to treated groups T_1_ (4% FS), T_2_ (6% FS), T_3_ (8% FS) as well as control group (T_0_) was evaluated and compared [15, 30].

### Statistical Analysis

The data obtained from testing was analyzed by IBM SPSS 23. 2-way ANOVA was applied to give the contrast of the average values of different parameters of three biochemical tests such as serum lipid profile test, ELISA and CBC of all the 4 groups (T_0_, T_1_, T_2_ and T_3_) after 15^th^, 30^th^ and 45^th^ days individually. Mean and Standard Error Mean (Mean±SEM) values were determined from the data and Post Hoc Tuckey’s test was applied to compare significant differences among individual treatment and control group. The difference between average values were considered significant in case of p≤0.05.

## Results

### Effect of flaxseeds on hematological parameters

In this study, the effect of flaxseed’s administration (4%, 6% and 8% from Day 0 to 45^th^ Day) on hematological parameters in rabbits were investigated. After the consumption of different doses of flaxseeds, the hematological parameters showed significant amendments (p≤0.05) in all treated groups, T_1_ (fed with 4% flaxseeds), T_2_ (6% flaxseeds) and T_3_ (8% flaxseeds) as compared to the control group. At concentration 8% of flaxseeds, the mean values of Hb (17.38±.00 mg/dL), RBCs (5.49±.05 ×10^6^/µL), HCT (47.97±.01 %), MCV (66.56±.00 fL), MCH (20.94±.00 pg), MCHC (46.59±.00 %), Platelets Count (664.18±3.68 ×10^3^/µL) and WBCs (11.66±.00 ×10^3^/µL) depicted maximum increase comparable to the control group (Table-1). The mean values of differential leukocytes neutrophils (53.95±.00), lymphocytes (45.95±.02), eosinophils (3.49±.00), basophils (7.05±.02^a^) and monocytes (12.00±.00) increased significantly (p≤0.05) when 4%, 6% and 8% of flaxseeds were given to the rabbits for consecutive 45 days (Table-2).

**Table 1.**
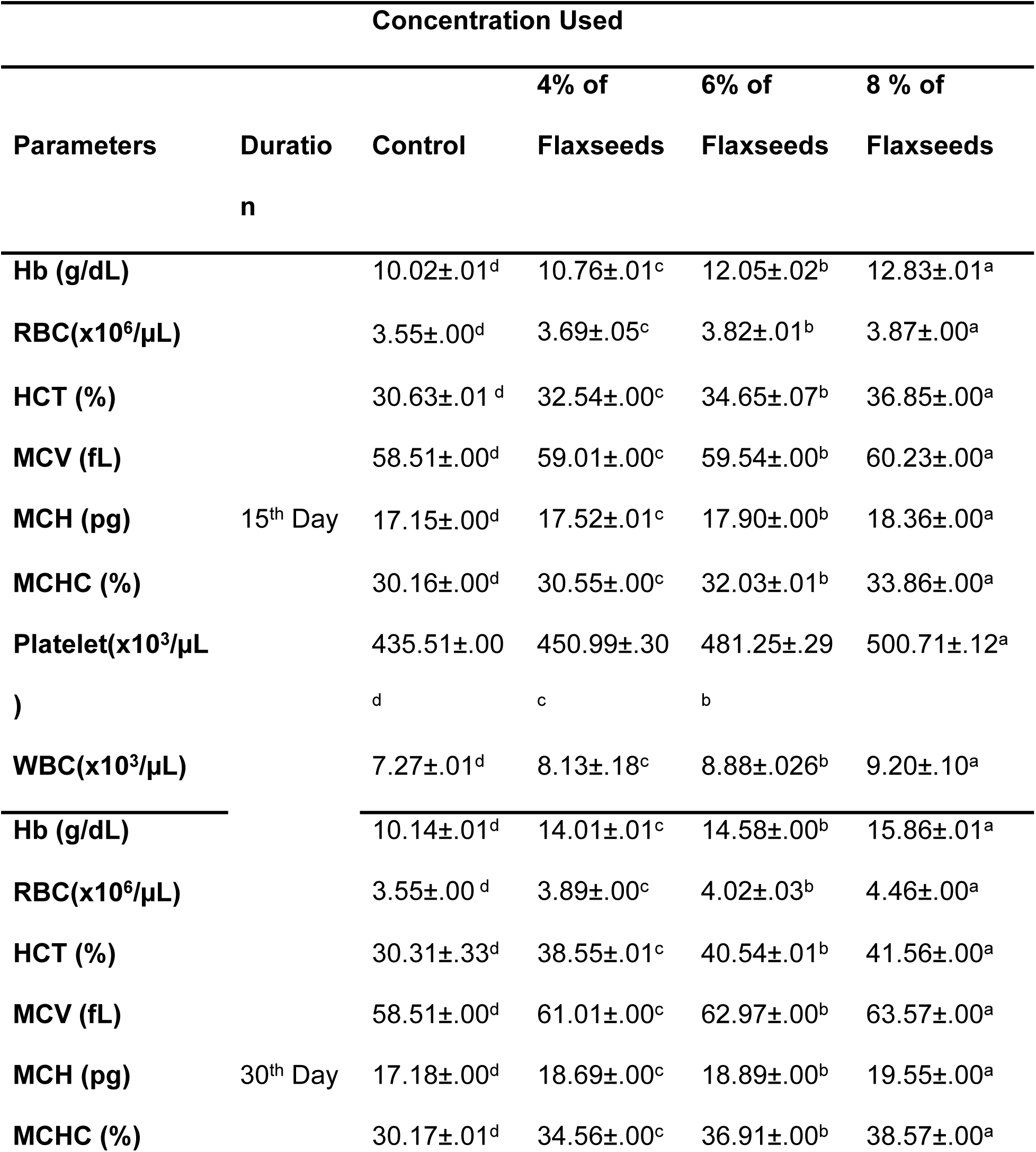
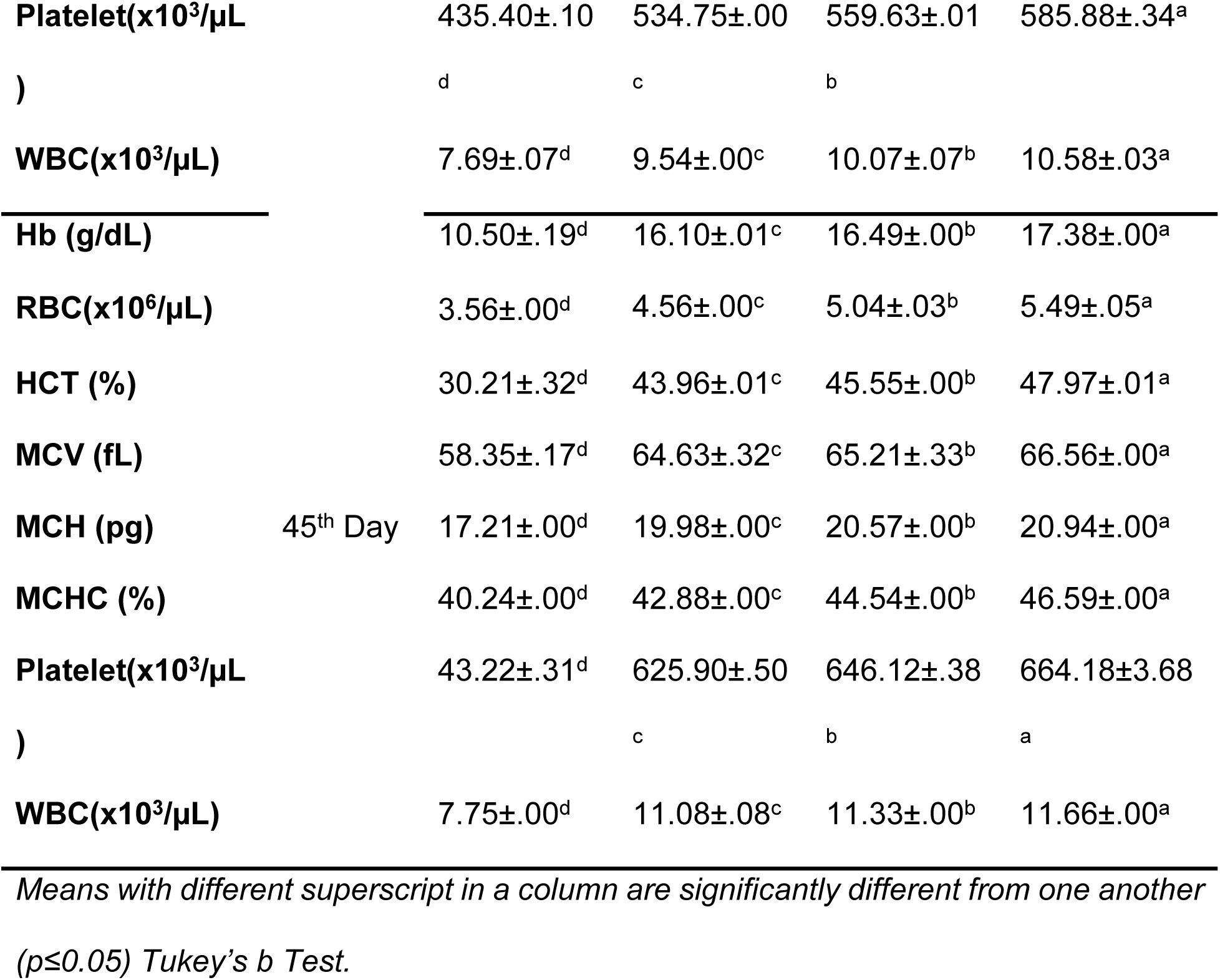
Effect of flaxseeds on hematological parameters of the rabbits at 15^th^, 30^th^ and 45^th^ day.

**Table 2.** Maximum effect of flaxseeds on differential leukocytes of the rabbits at following concentrations at 15^th^, 30^th^ and 45^th^ day (Mean±S.E.M)

| Parameters | Duration | Concentration Used |  |  |  |
| --- | --- | --- | --- | --- | --- |
|  |  | Control | 4% of Flaxseeds | 6% of Flaxseeds | 8 % of Flaxseeds |
| **Neutrophils (%)** | 15<sup>th</sup> Day | 37.03±.00<sup>d</sup> | 38.00±.00<sup>c</sup> | 38.66±.00<sup>b</sup> | 39.87±.00<sup>a</sup> |
| **Lymphocytes(%)** |  | 28.01±.00<sup>d</sup> | 28.56±.00<sup>c</sup> | 30.65±.00<sup>b</sup> | 32.78±.00<sup>a</sup> |
| **Eosinophils (%)** |  | .51±.00<sup>d</sup> | .98±.00<sup>c</sup> | 1.56±.00<sup>b</sup> | 1.96±.00<sup>a</sup> |
| **Basophils (%)** |  | 2.49±.00<sup>d</sup> | 3.01±.00<sup>c</sup> | 3.52±.01<sup>b</sup> | 3.90±.00<sup>a</sup> |
| **Monocytes (%)** |  | 4.01±.00<sup>d</sup> | 5.04±.00<sup>c</sup> | 5.68±.00<sup>b</sup> | 6.54±.00<sup>a</sup> |
| **Neutrophils (%)** | 30<sup>th</sup> Day | 37.03±.00<sup>d</sup> | 41.55±.00<sup>c</sup> | 43.43±.00<sup>b</sup> | 45.16±.00<sup>a</sup> |
| **Lymphocytes(%)** |  | 28.01±.00<sup>d</sup> | 34.28±.00<sup>c</sup> | 35.96±.00<sup>b</sup> | 37.20±.00<sup>a</sup> |
| **Eosinophils (%)** |  | .51±.01<sup>d</sup> | 2.27±.00<sup>c</sup> | 2.52±.00<sup>b</sup> | 2.87±.00<sup>a</sup> |
| **Basophils (%)** |  | 2.49±.00<sup>d</sup> | 4.42±.00<sup>c</sup> | 4.87±.00<sup>b</sup> | 5.25±.00<sup>a</sup> |
| **Monocytes (%)** |  | 4.01±.00<sup>d</sup> | 7.34±.00<sup>c</sup> | 8.55±.00<sup>b</sup> | 9.30±.00<sup>a</sup> |
| **Neutrophils (%)** | 45<sup>th</sup> Day | 37.03±.00<sup>d</sup> | 47.41±.04<sup>c</sup> | 50.33±.00<sup>b</sup> | 53.95±.00<sup>a</sup> |
| **Lymphocytes(%)** |  | 26.35±.66<sup>d</sup> | 40.70±.00<sup>c</sup> | 43.88±.00<sup>b</sup> | 45.95±.02<sup>a</sup> |
| **Eosinophils (%)** |  | .53±.00<sup>d</sup> | 3.06±.00<sup>c</sup> | 3.31±.00<sup>b</sup> | 3.49±.00<sup>a</sup> |
| **Basophils (%)** |  | 2.48±.00<sup>d</sup> | 5.88±.00<sup>c</sup> | 6.51±.00<sup>b</sup> | 7.05±.02<sup>a</sup> |
| **Monocytes (%)** |  | 4.02±.01<sup>d</sup> | 10.45±.00<sup>c</sup> | 11.66±.00<sup>b</sup> | 12.00±.00<sup>a</sup> |

*Means with different superscript in a column are significantly different from one another (p≤0<sup>a</sup>.05) Tukey's b Test.*

Maximum increase in the levels of all hematological parameters obtained in group T_3_ treated with high dose = 8% flaxseeds, followed by group T_2_ (6% flaxseeds) and group T_1_ (4% flaxseeds) (Figure-1-13).

**Figure 1.**
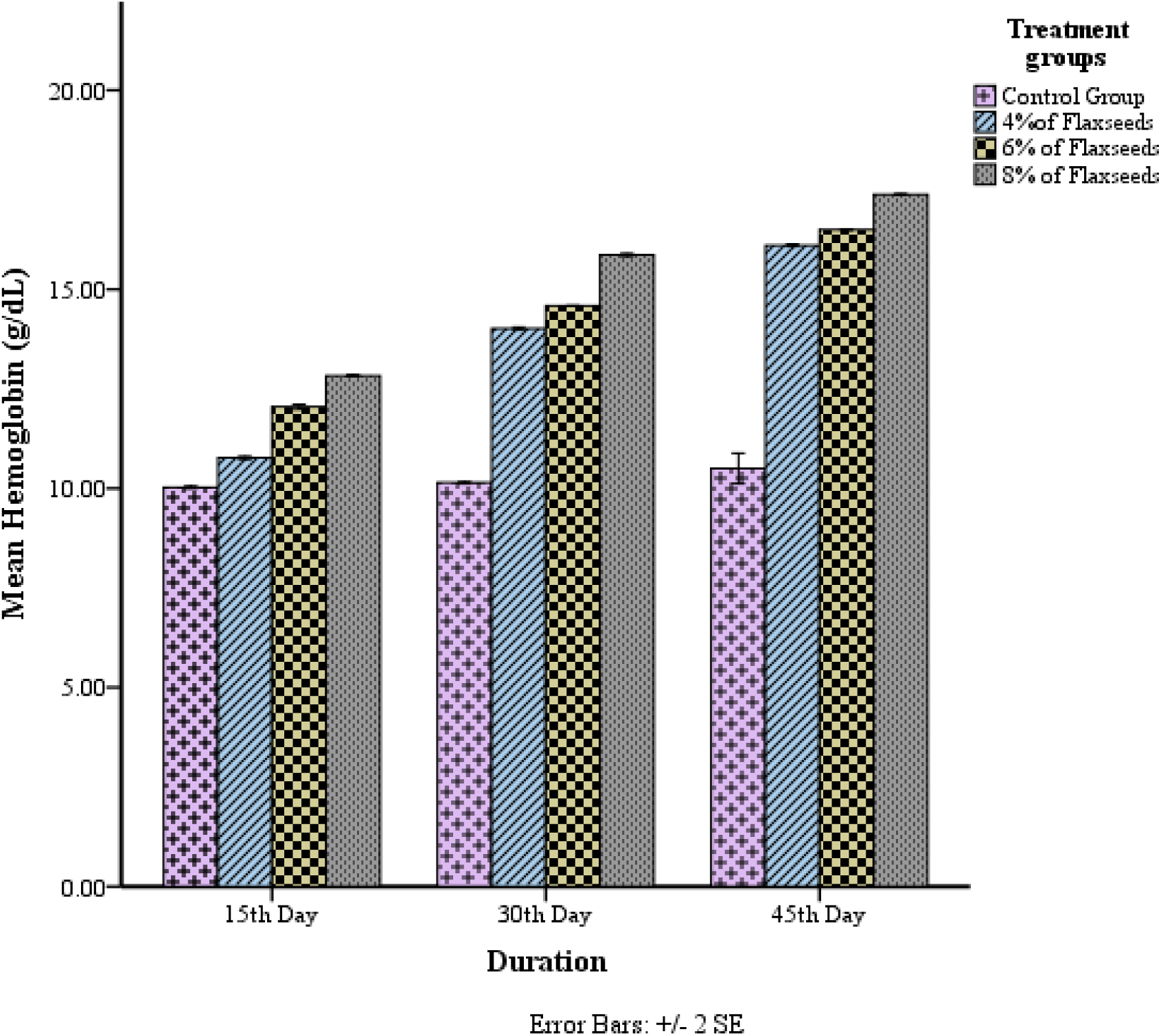
Concentration and time dependent effect of flaxseeds on hemoglobin (Hb) level (g/dL) of rabbits in T_0_ (Control), T_1_ (4% flaxseeds), T_2_ (6% flaxseeds) and T_3_ (8% flaxseeds) groups at different time-scales (15th, 30th and 45th days).

**Figure 2.**
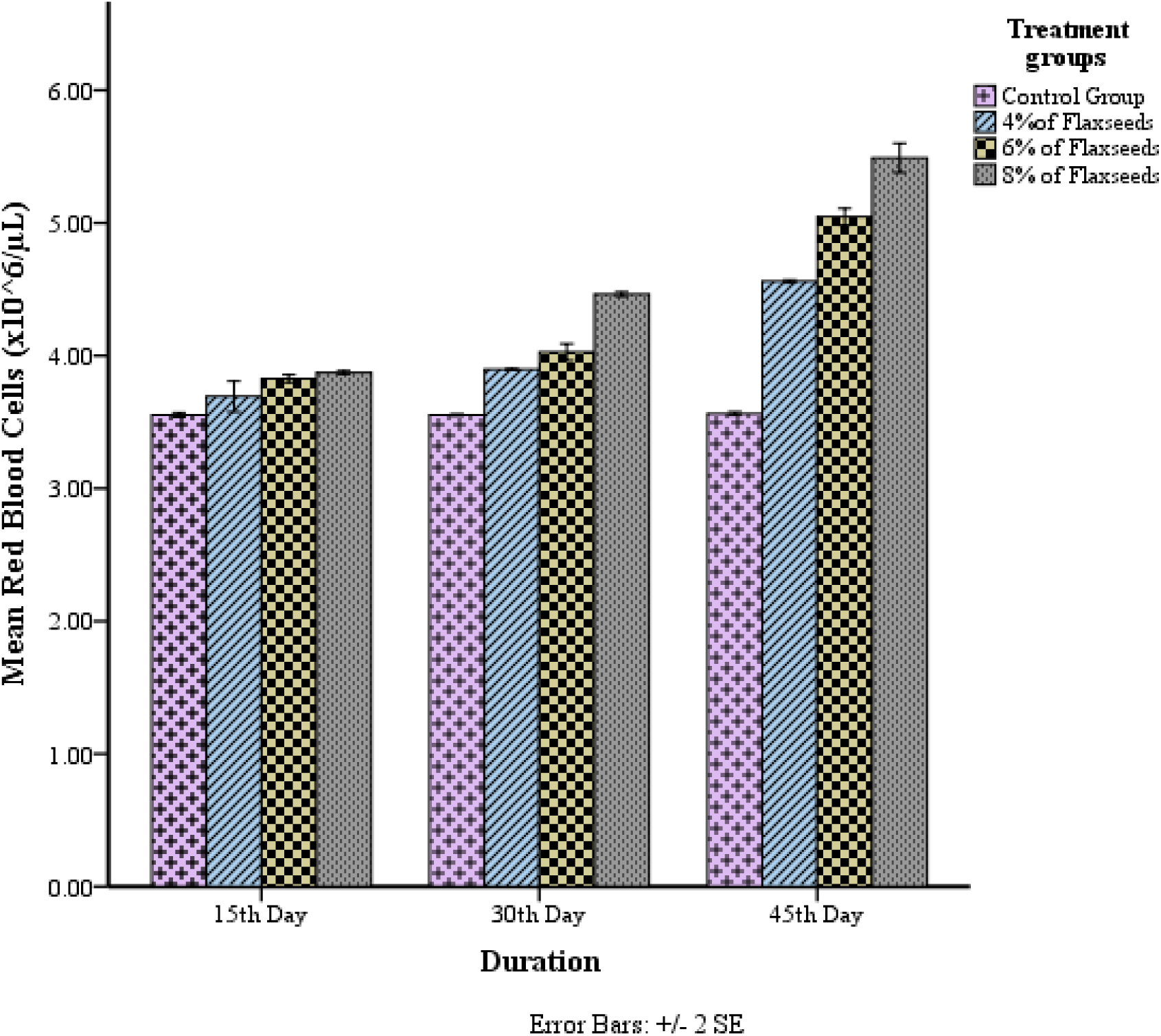
Concentration and time dependent effect of flaxseeds on red blood cells (RBCs) (×10^6^ /µL)) of rabbits in T_0_ (Control), T_1_ (4% flaxseeds), T_2_ (6% flaxseeds) and T_3_ (8% flaxseeds) groups at different time-scales (15th, 30th and 45th days).

**Figure 3.**
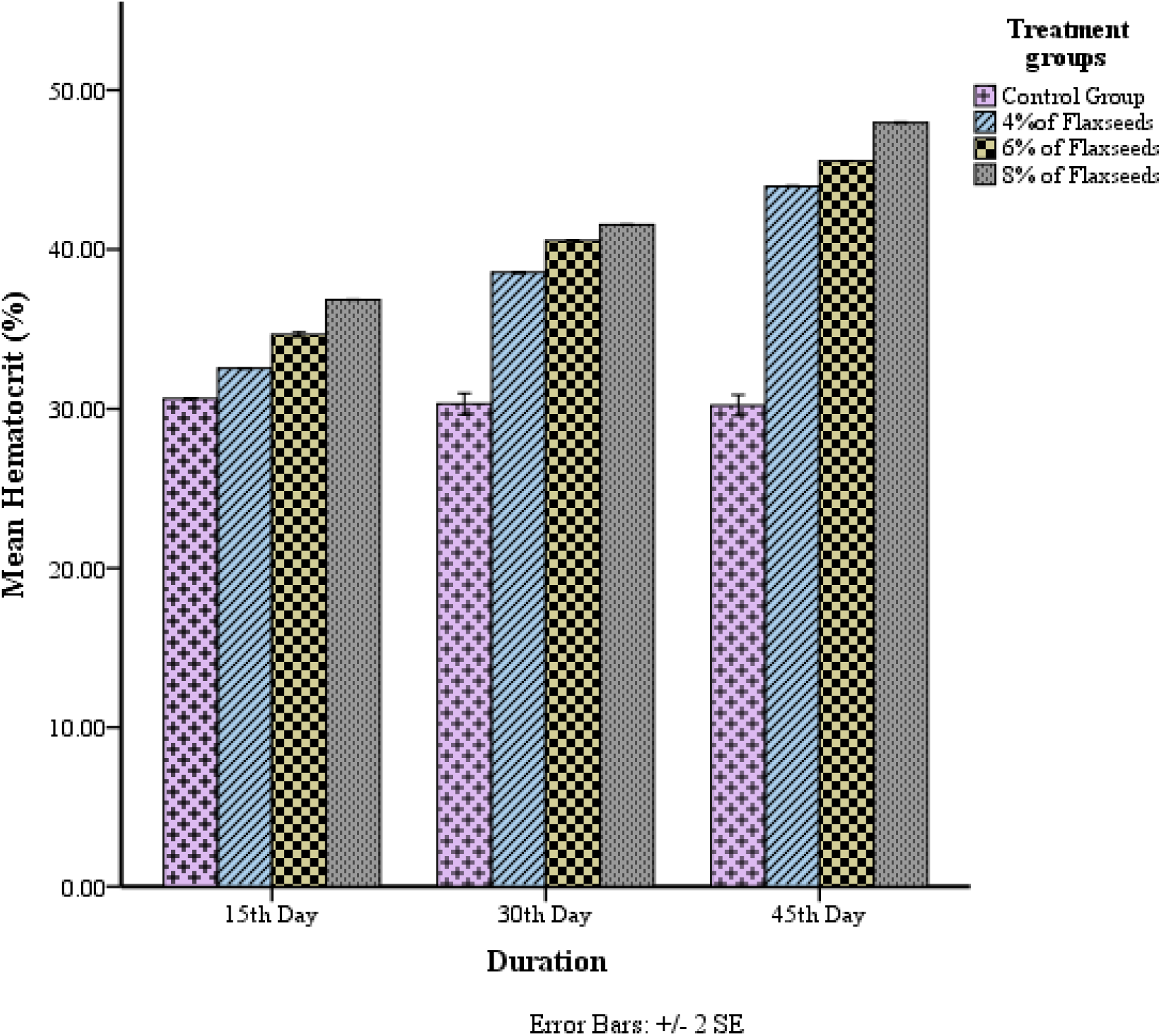
Concentration and time dependent effect of flaxseeds on hematocrit (%) of rabbits in T_0_ (Control), T_1_ (4% flaxseeds), T_2_ (6% flaxseeds) and T_3_ (8% flaxseeds) groups at different time-scales (15th, 30th and 45th days).

**Figure 4.**
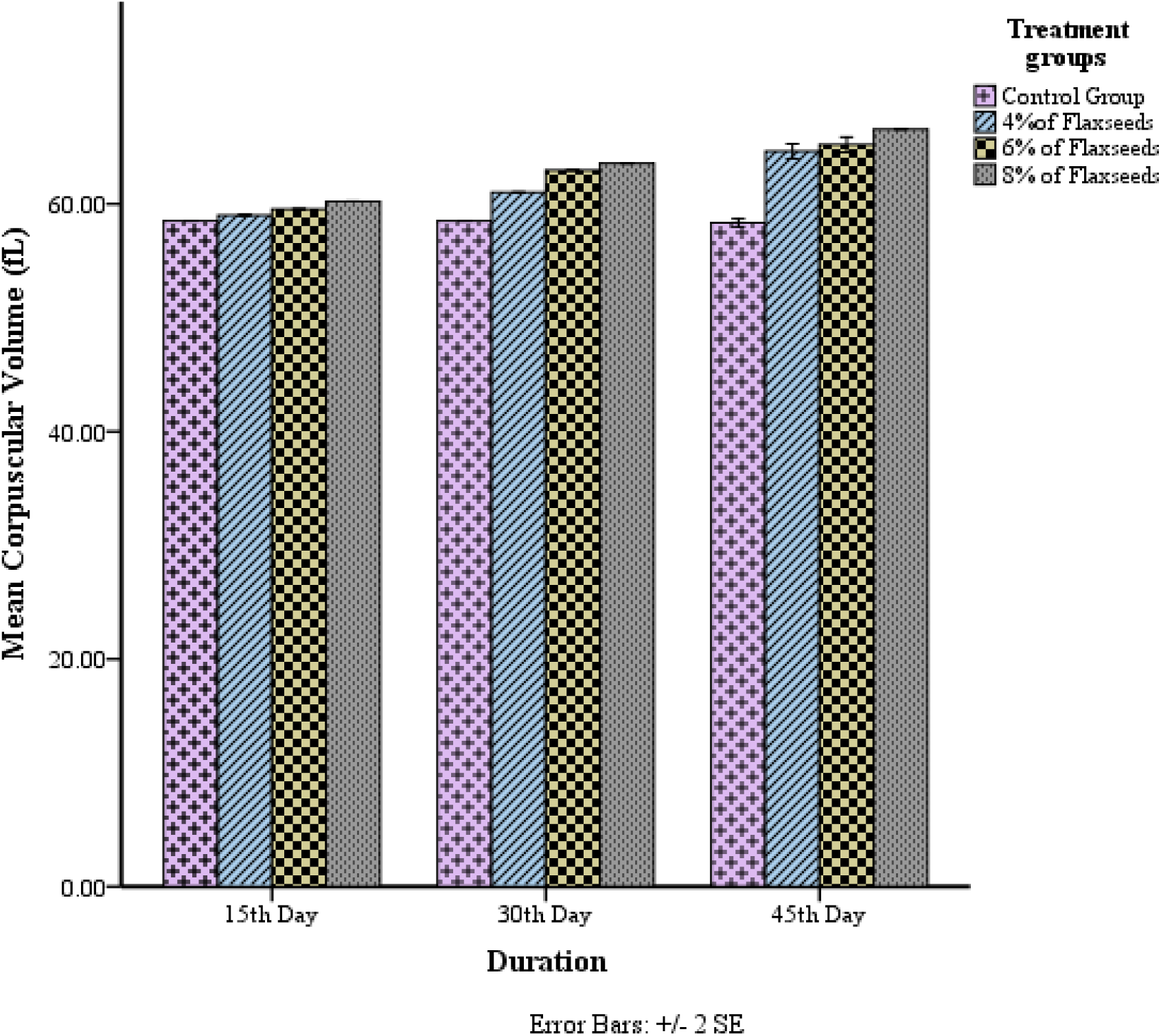
Concentration and time dependent effect of flaxseeds on MCV (fL) level of rabbits in T_0_ (Control), T_1_ (4% flaxseeds), T_2_ (6% flaxseeds) and T_3_ (8% flaxseeds) groups at different time-scales (15th, 30th and 45th days).

**Figure 5.**
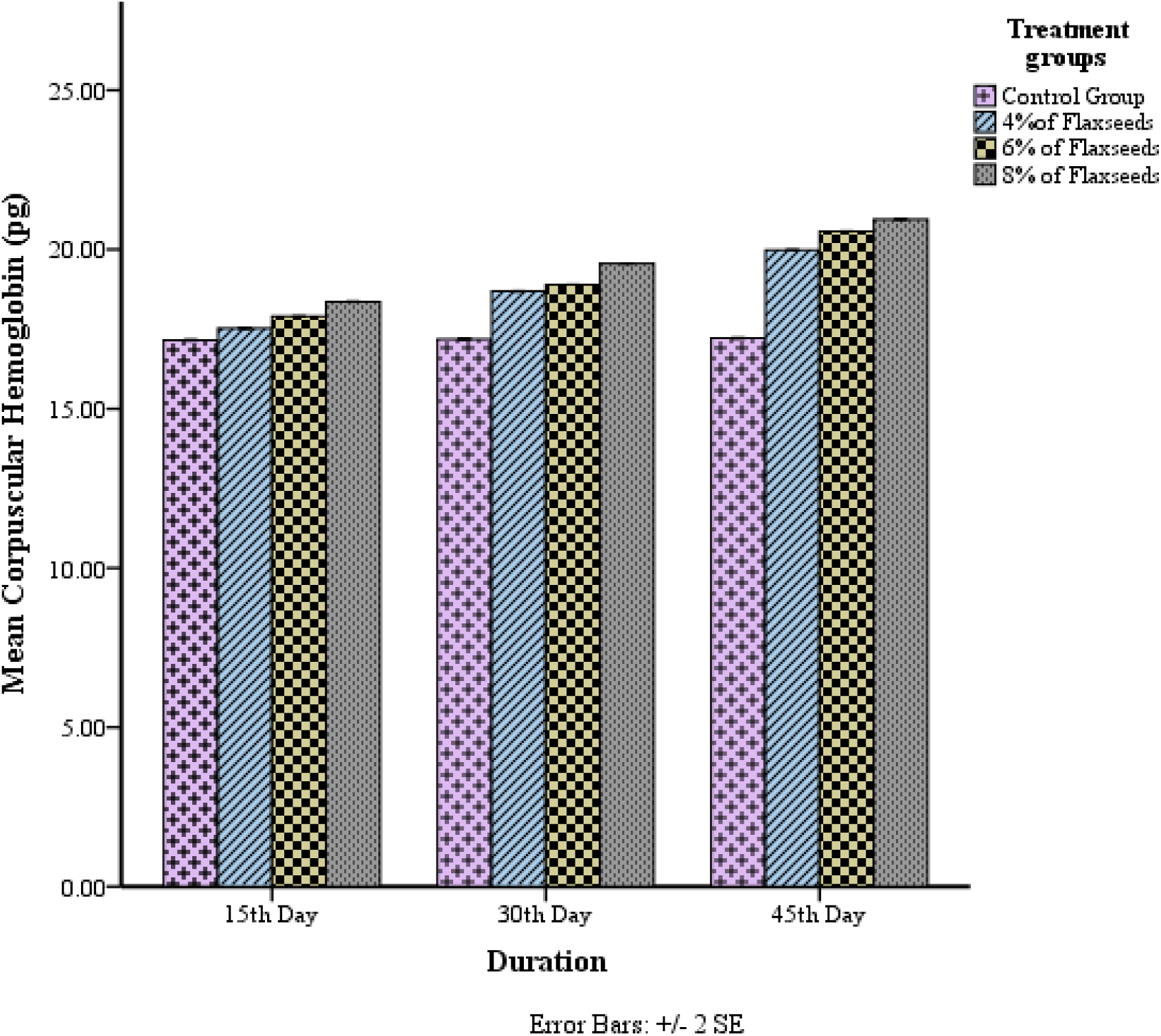
Concentration and time dependent effect of flaxseeds on MCH (pg) level of rabbits in T_0_ (Control), T_1_ (4% flaxseeds), T_2_ (6% flaxseeds) and T_3_ (8% flaxseeds) groups at different time-scales (15th, 30th and 45th days).

**Figure 6.**
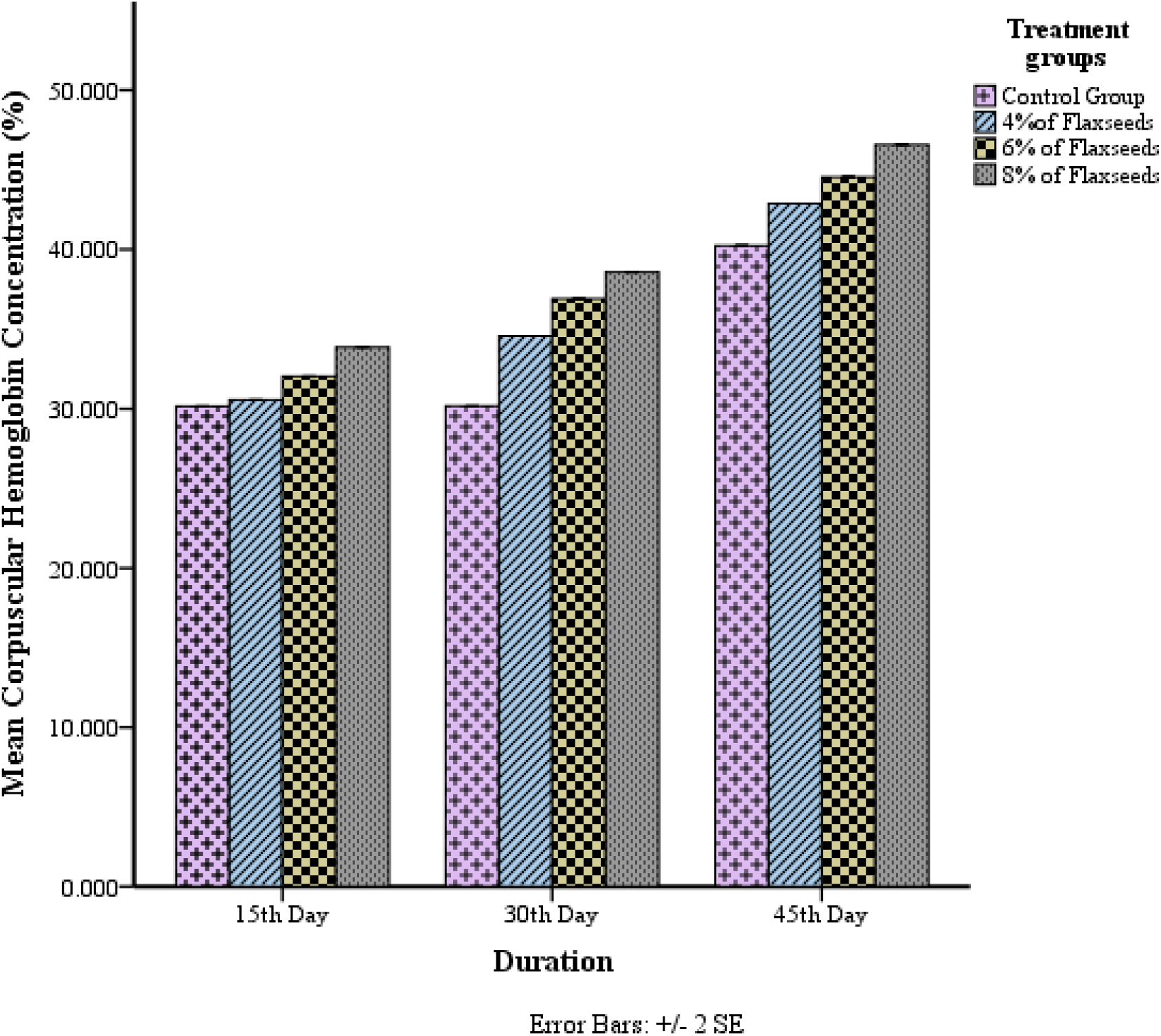
Concentration and time dependent effect of flaxseeds on MCHC (%) level of rabbits in T_0_ (Control), T_1_ (4% flaxseeds), T_2_ (6% flaxseeds) and T_3_ (8% flaxseeds) groups at different time-scales (15th, 30th and 45th days).

**Figure 7.**
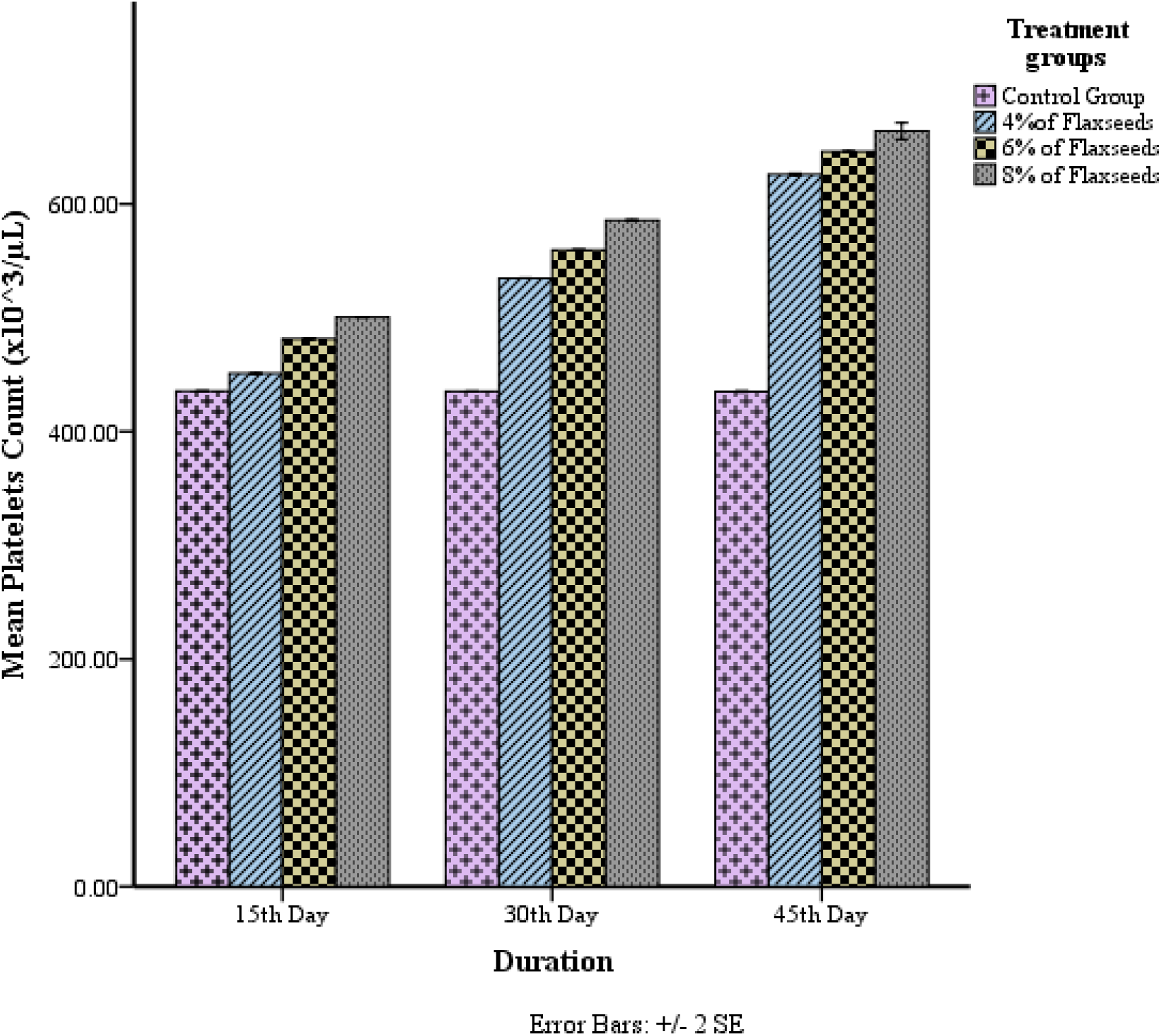
Concentration and time dependent effect of flaxseeds on platelet counts (×10^3^ /µL) of rabbits in T_0_ (Control), T_1_ (4% flaxseeds), T_2_ (6% flaxseeds) and T_3_ (8% flaxseeds) groups at different time-scales (15th, 30th and 45th days).

**Figure 8.**
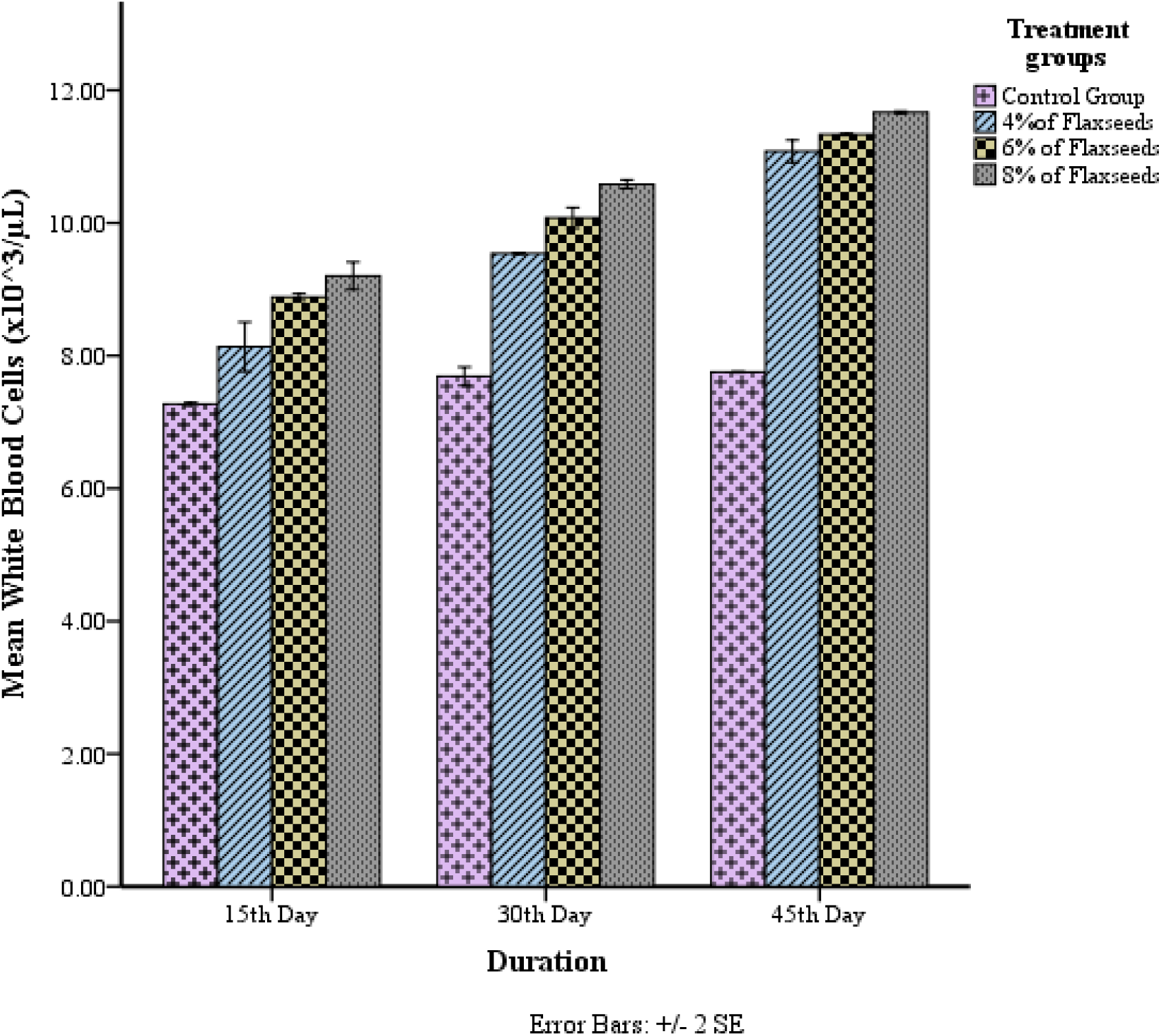
Concentration and time dependent effect of flaxseeds on white blood cells (WBCs) (×10^3^ /µL) of rabbits in T_0_ (Control), T_1_ (4% flaxseeds), T_2_ (6% flaxseeds) and T_3_ (8% flaxseeds) groups at different time-scales (15th, 30th and 45th days).

**Figure 9.**
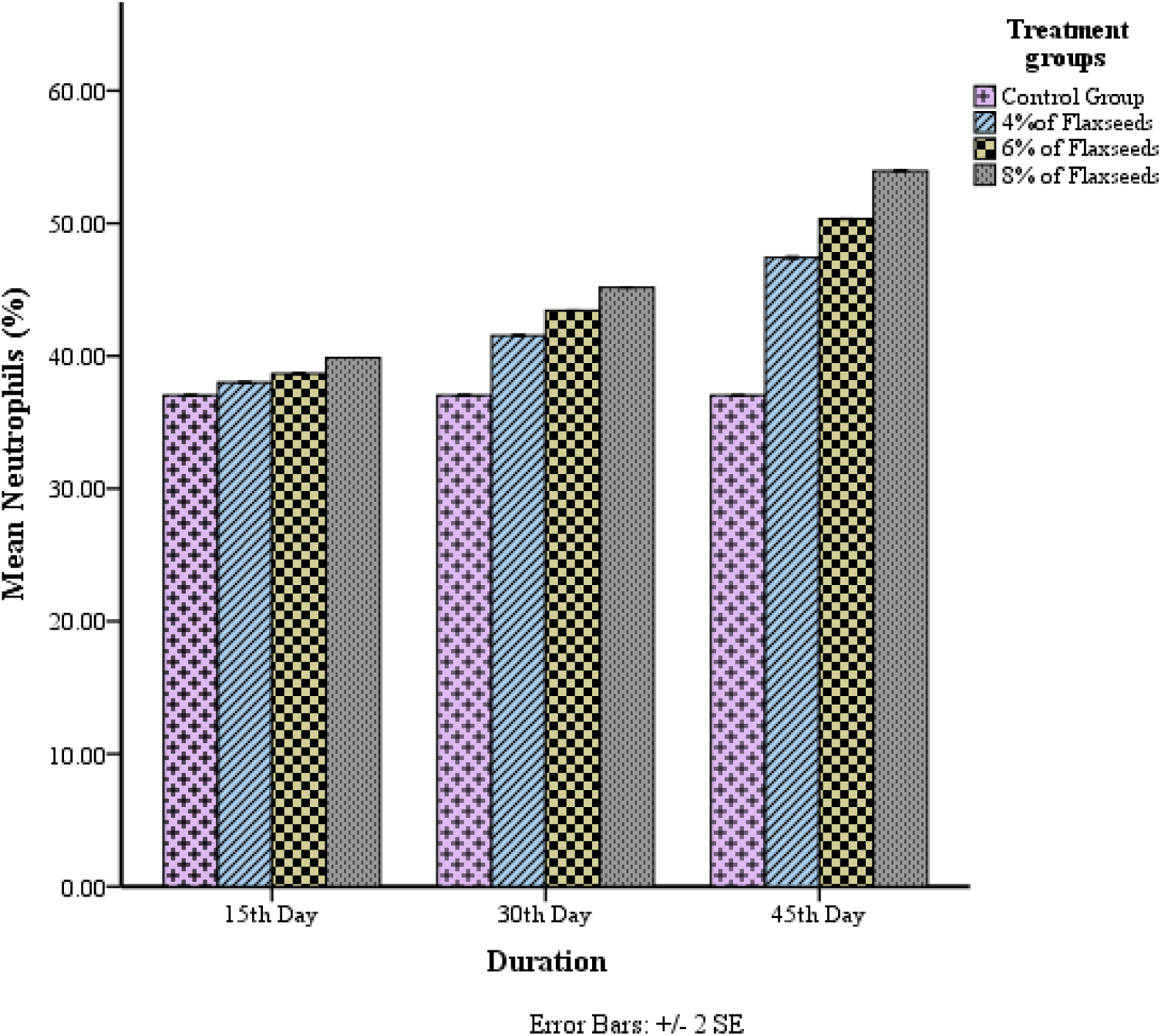
Concentration and time dependent effect of flaxseeds on neutrophils (%) level of rabbits in T_0_ (Control), T_1_ (4% flaxseeds), T_2_ (6% flaxseeds) and T_3_ (8% flaxseeds) groups at different time-scales (15th, 30th and 45th days.

**Figure 10.**
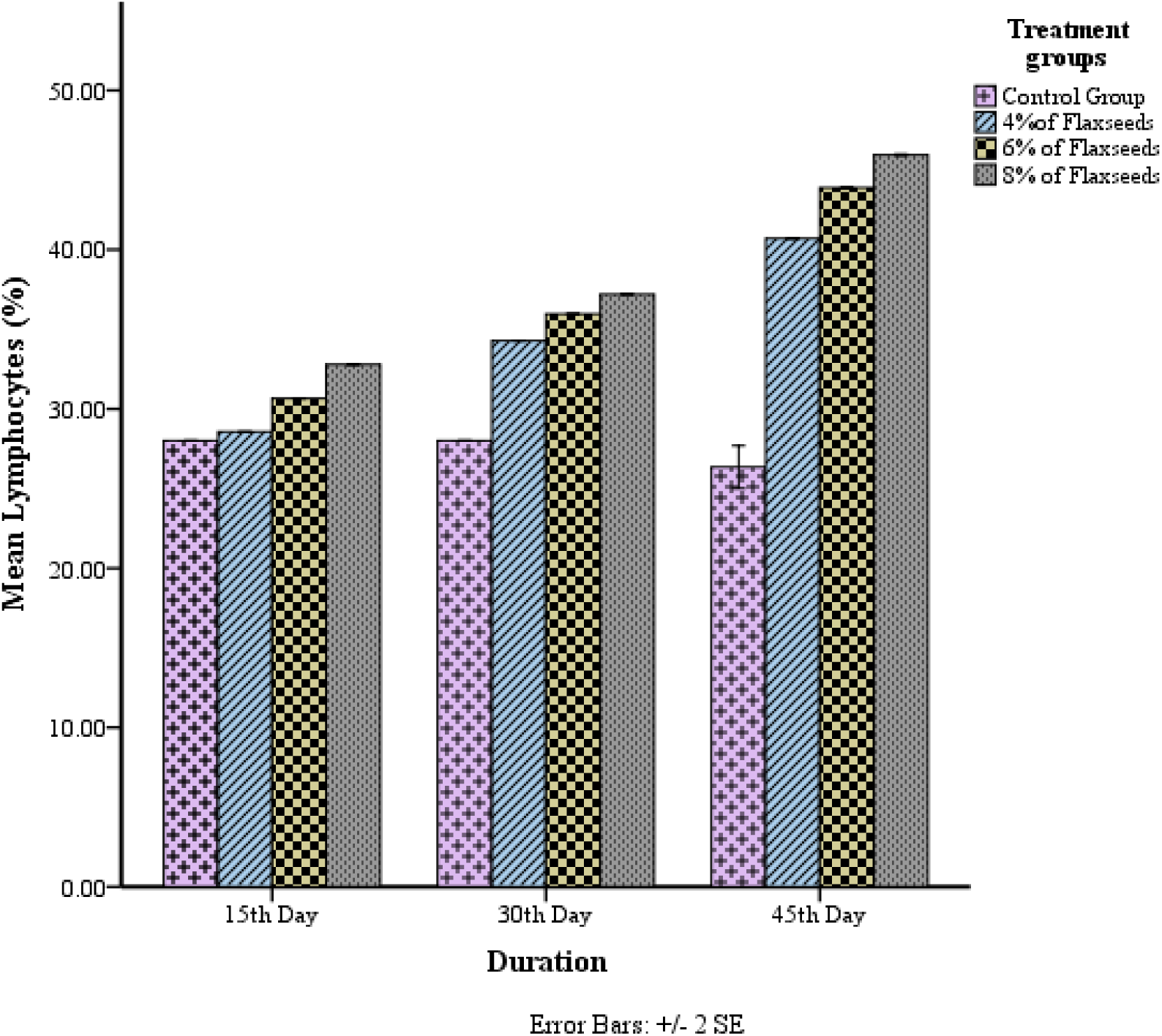
Concentration and time dependent effect of flaxseeds on lymphocytes (%) level of rabbits in T_0_ (Control), T_1_ (4% flaxseeds), T_2_ (6% flaxseeds) and T_3_ (8% flaxseeds) groups at different time-scales (15th, 30th and 45th days).

**Figure 11.**
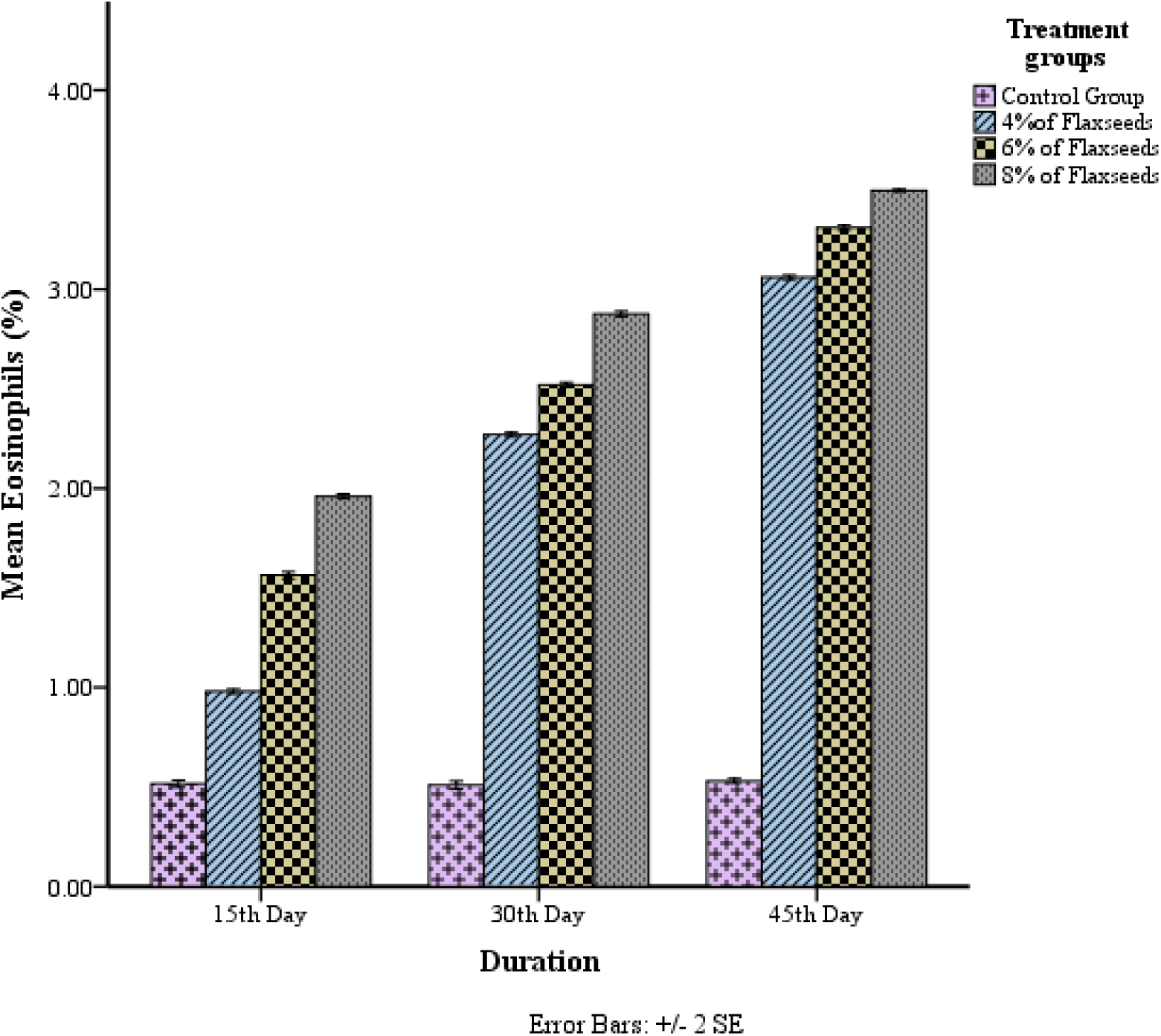
Concentration and time dependent effect of flaxseeds on eosinophils (%) level of rabbits in T_0_ (Control), T_1_ (4% flaxseeds), T_2_ (6% flaxseeds) and T_3_ (8% flaxseeds) groups at different time-scales (15th, 30th and 45th days).

**Figure 12.**
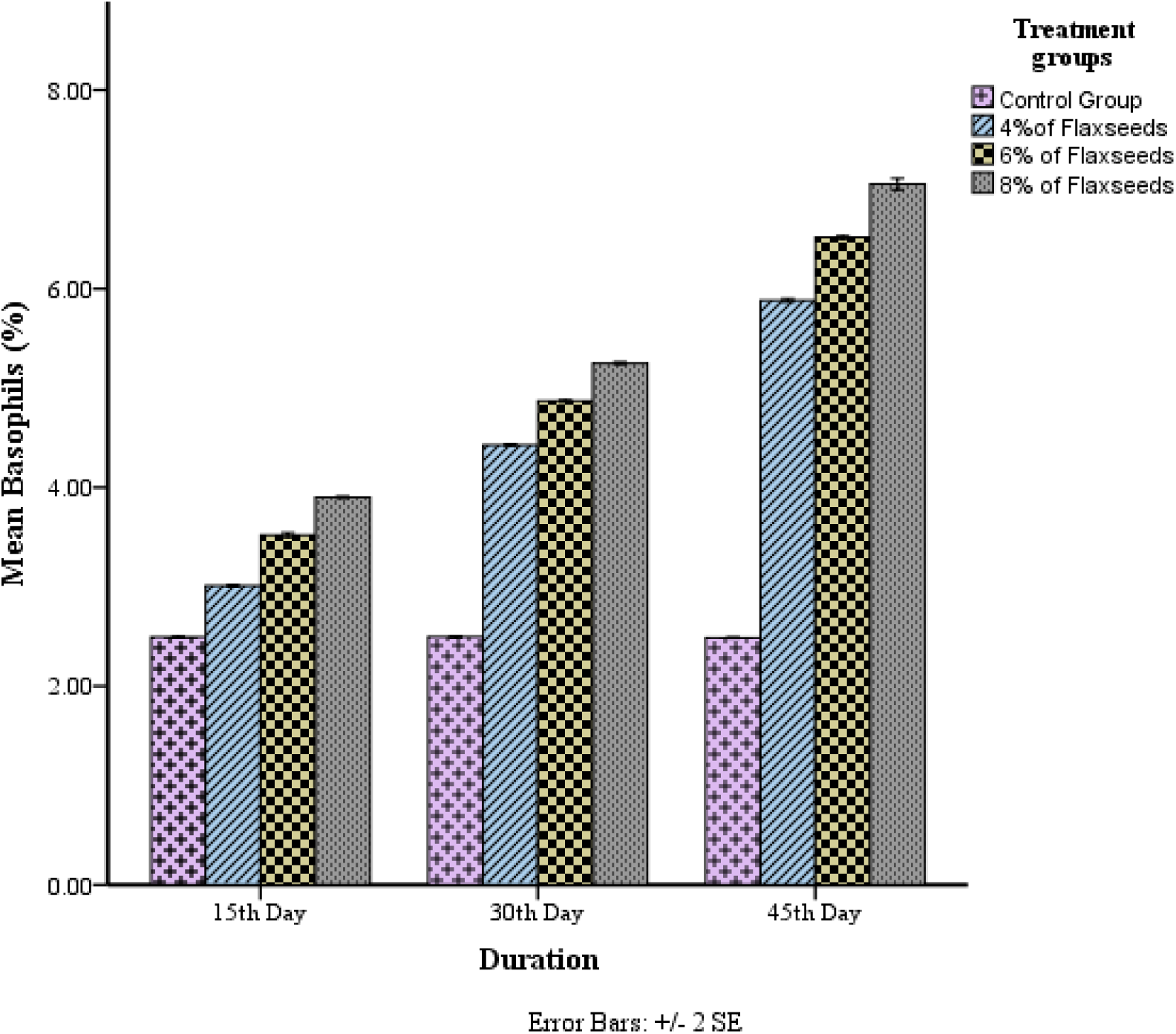
Concentration and time dependent effect of flaxseeds on basophils (%) level of rabbits in T_0_ (Control), T_1_ (4% flaxseeds), T_2_ (6% flaxseeds) and T_3_ (8% flaxseeds) groups at different time-scales (15th, 30th and 45th days).

**Figure 13.**
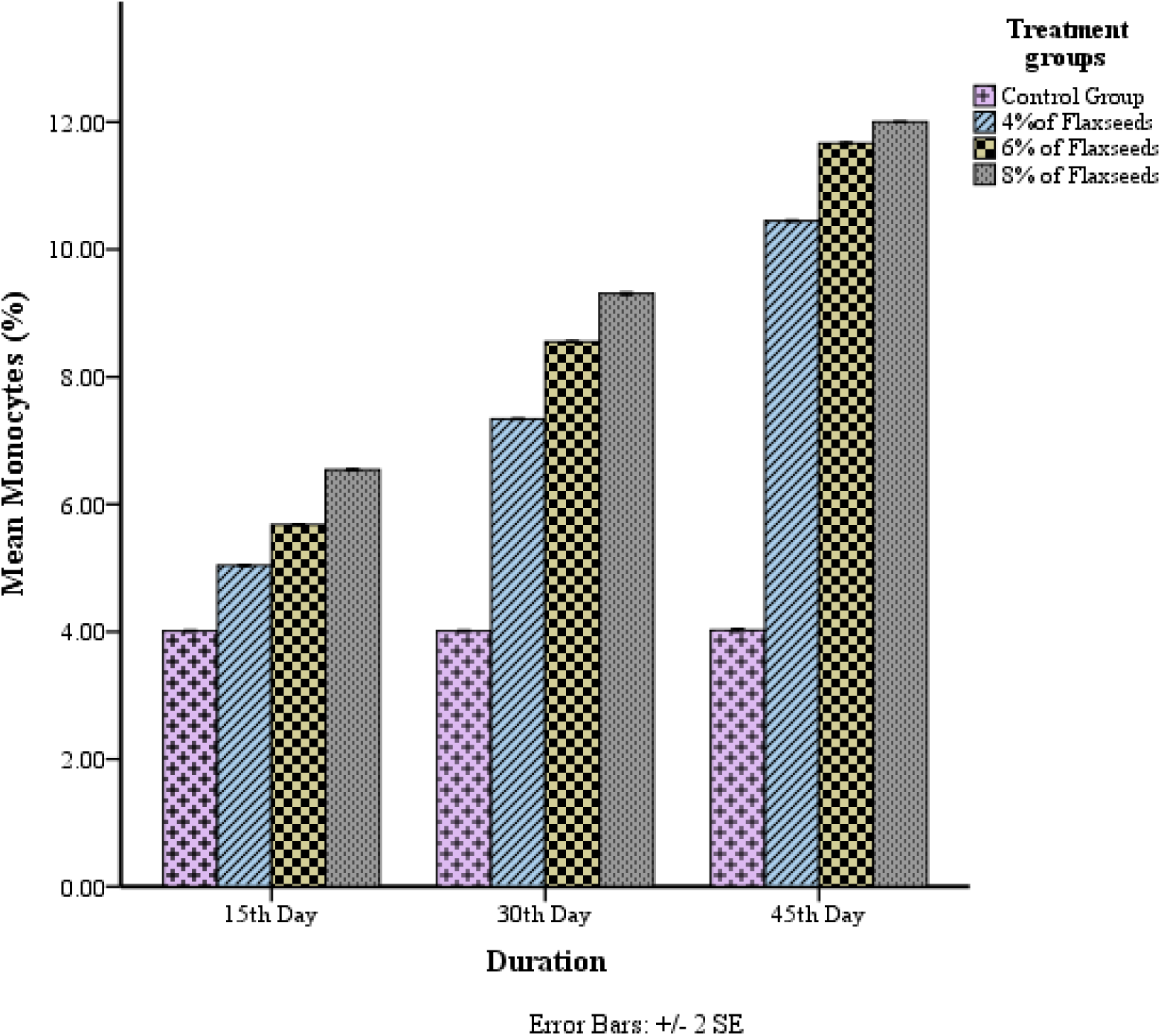
Concentration and time dependent effect of flaxseeds on monocytes (%) level of rabbits in T_0_ (Control), T_1_ (4% flaxseeds), T_2_ (6% flaxseeds) and T_3_ (8% flaxseeds) groups at different time-scales (15th, 30th and 45th days).

### Effect of flaxseeds on lipid profile

The lipid profile of the male rabbits restored significantly (p≤0.05) after the consumption of different doses of flaxseeds as compared to the control group. Flaxseeds consumption resulted significant changes in the mean values of total cholesterol (53.3300±.0 mg/dL), serum triglycerides (17.64±.00 mg/dL), and low-density lipoprotein (22.57±.01 mg/dL) and high-density lipoprotein (47.27±.01 mg/dL) at concentration 8% of flaxseeds comparable to the control group (Table-3). Significant reduction in total cholesterol (TC), serum triglycerides (TG), and low-density lipoprotein (LDL-C) levels and significant increase in high-density lipoprotein (HDL-C) were recorded in group T_3_ treated with high dose = 8% flaxseeds, followed by group T_2_ (6% flaxseeds) and group T_1_ (4% flaxseeds), summarized in (Figure-14-17).

**Table 3.** Maximum effect of flaxseeds on lipid profile of the rabbits at following concentrations 15^th^, 30^th^ and 45^th^ day (Mean±S.E.M).

| Parameters | Duration | Concentration Used |  |  |  |
| --- | --- | --- | --- | --- | --- |
|  |  | Control | 4% of<br>Flaxseeds | 6% of<br>Flaxseeds | 8 % of<br>Flaxseeds |
| TC (mg/dL) | 15<sup>th</sup> Day | 76.37±.01<sup>d</sup> | 73.05±.02<sup>c</sup> | 70.74±.00<sup>b</sup> | 68.15±.01<sup>a</sup> |
| TAG (mg/dL) |  | 44.55±.01<sup>d</sup> | 40.95±.02<sup>c</sup> | 37.09±.00<sup>b</sup> | 25.65±.01<sup>a</sup> |
| LDL-C (mg/dL) |  | 50.49±.00<sup>d</sup> | 47.24±.01<sup>c</sup> | 44.10±.00<sup>b</sup> | 41.64±.01<sup>a</sup> |
| HDL-C<br>(mg/dL) |  | 17.63±.00<sup>d</sup> | 20.45±.00<sup>c</sup> | 24.08±.00<sup>b</sup> | 27.75±.01<sup>a</sup> |
| TC (mg/dL) | 30<sup>th</sup> Day | 76.07±.33<sup>d</sup> | 65.22±.01<sup>c</sup> | 63.03±.03<sup>b</sup> | 60.57±.00<sup>a</sup> |
| TAG (mg/dL) |  | 44.03±.02<sup>d</sup> | 31.49±.00<sup>c</sup> | 28.03±.01<sup>b</sup> | 25.65±.01<sup>a</sup> |
| LDL-C (mg/dL) |  | 50.53±.01<sup>d</sup> | 38.32±.01<sup>c</sup> | 35.00±.00<sup>b</sup> | 32.65±.00<sup>a</sup> |
| HDL-C<br>(mg/dL) |  | 17.65±.00<sup>d</sup> | 30.14±.00<sup>c</sup> | 33.65±.00<sup>b</sup> | 37.13±.01<sup>a</sup> |
| TC (mg/dL) | 45<sup>th</sup> Day | 76.48±.01<sup>d</sup> | 56.02±.00<sup>c</sup> | 53.24±.01<sup>b</sup> | 53.33±.01<sup>a</sup> |
| TAG (mg/dL) |  | 44.32±.00<sup>d</sup> | 21.22±.00<sup>c</sup> | 18.54±.00<sup>b</sup> | 17.64±.00<sup>a</sup> |
| LDL-C (mg/dL) |  | 50.56±.00<sup>d</sup> | 29.85±.00<sup>c</sup> | 26.20±.00<sup>b</sup> | 22.57±.01<sup>a</sup> |
| HDL-C<br>(mg/dL) |  | 17.66±.00<sup>d</sup> | 40.09±.01<sup>c</sup> | 43.46±.01<sup>b</sup> | 47.27±.01<sup>a</sup> |

*Means with different superscript in a column are significantly different from one another (p≤0.05) Tukey's b Test.*

**Figure 14.**
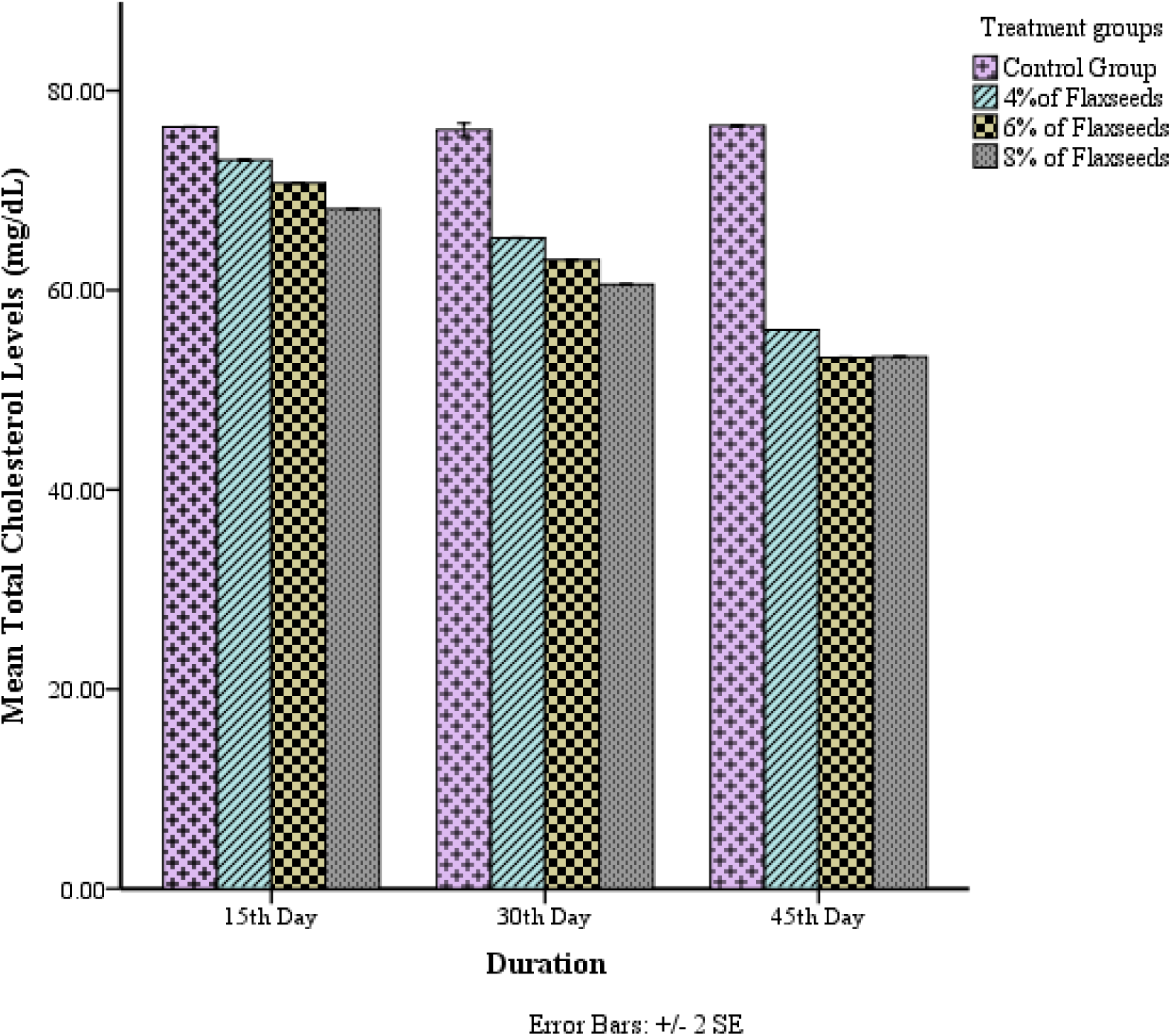
Concentration and time dependent effect of flaxseeds on total cholesterol level (mg/dL) of rabbits in T_0_ (Control), T_1_ (4% flaxseeds), T_2_ (6% flaxseeds) and T_3_ (8% flaxseeds) groups at different time-scales (15^th^, 30^th^ and 45^th^ days).

**Figure 15.**
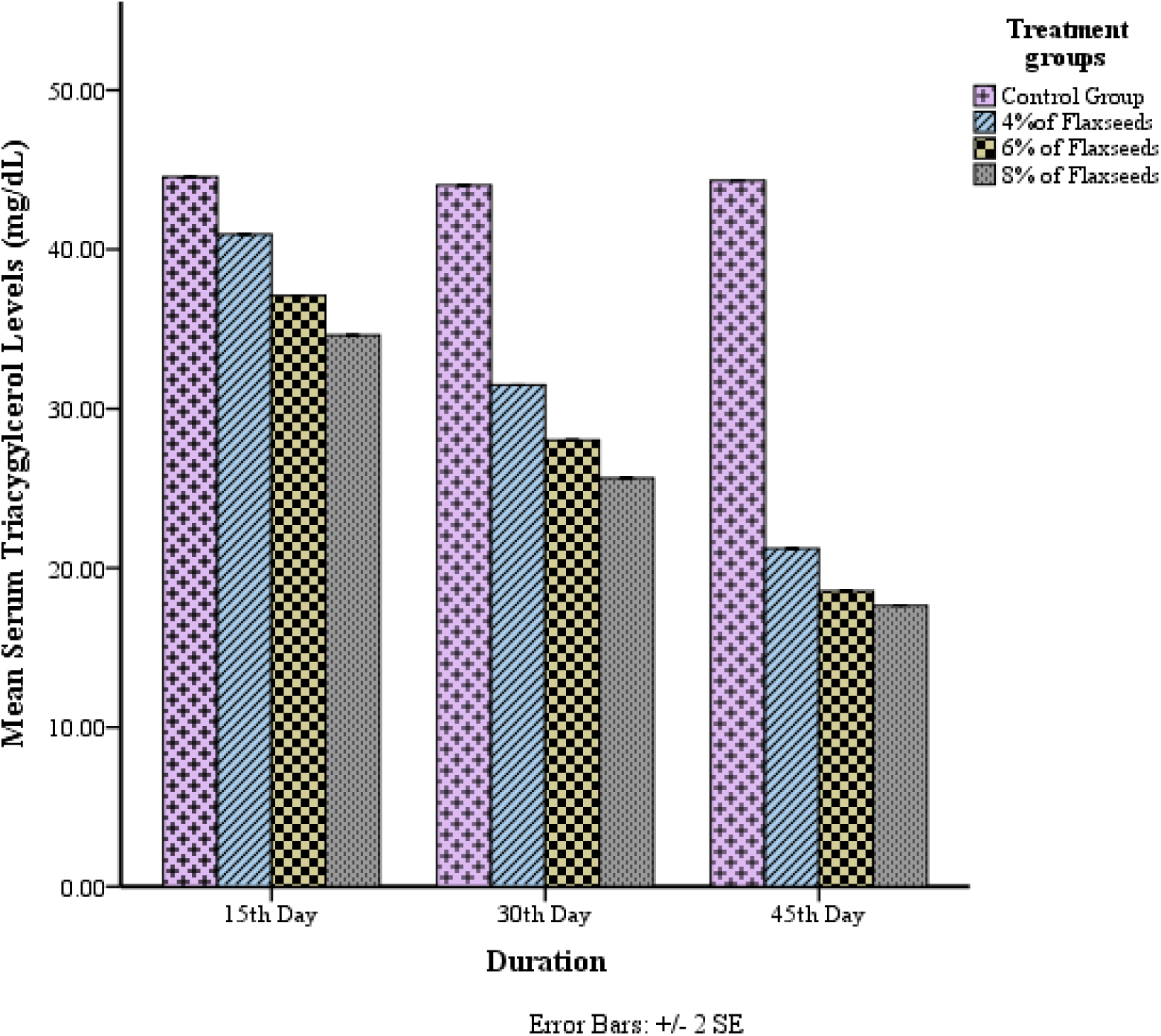
Concentration and time dependent effect of flaxseeds on serum triacylglycerol (TAG) level (mg/dL) of rabbits in T_0_ (Control), T_1_ (4% flaxseeds), T_2_ (6% flaxseeds) and T_3_ (8% flaxseeds) groups at different time-scales (15^th^, 30^th^ and 45^th^ days).

**Figure 16.**
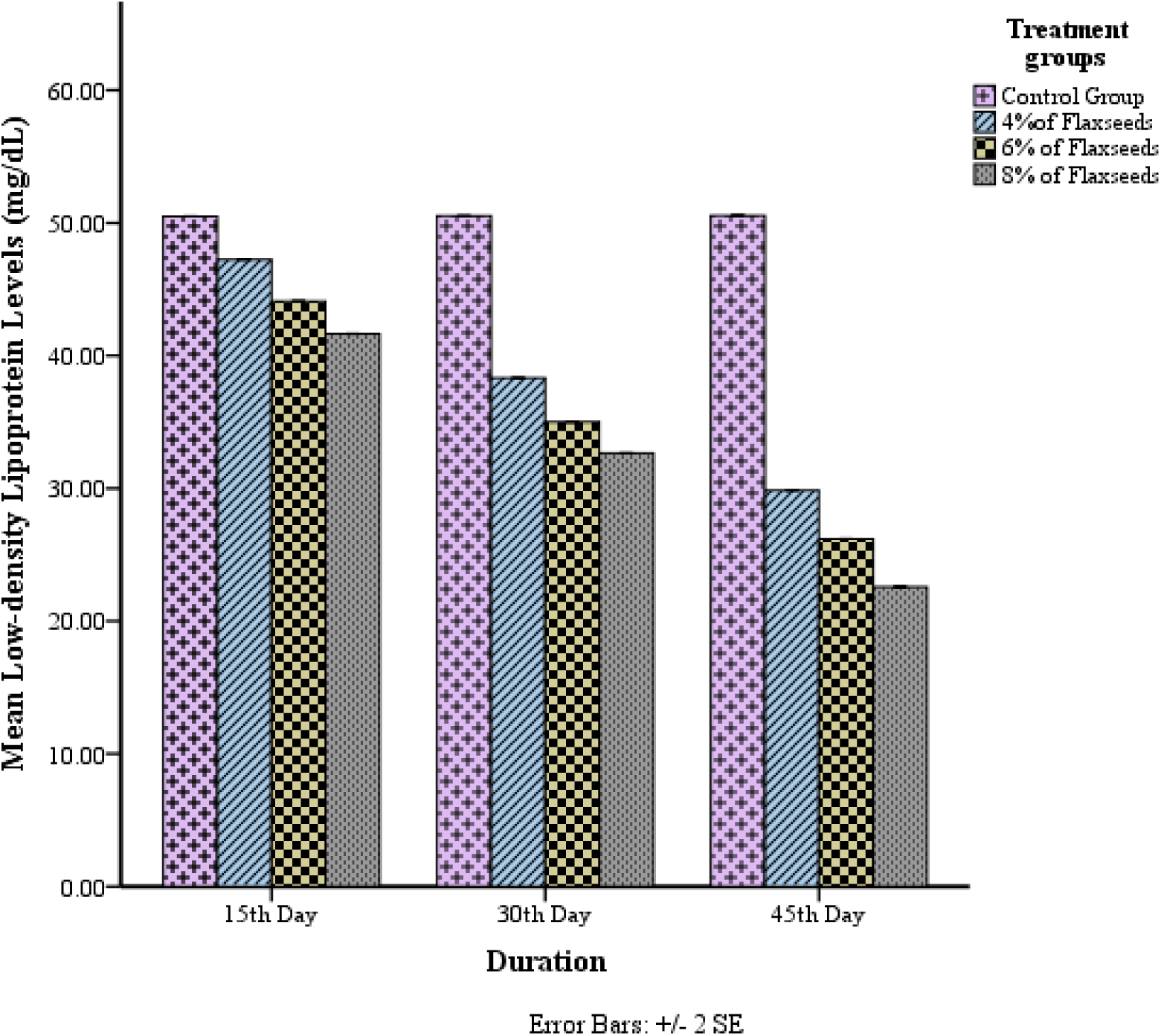
Concentration and time dependent effect of flaxseeds on low-density lipoprotein (LDL) level (mg/dL) of rabbits in T_0_ (Control), T_1_ (4% flaxseeds), T_2_ (6% flaxseeds) and T_3_ (8% flaxseeds) groups at different time-scales (15th, 30th and 45th days).

**Figure 17.**
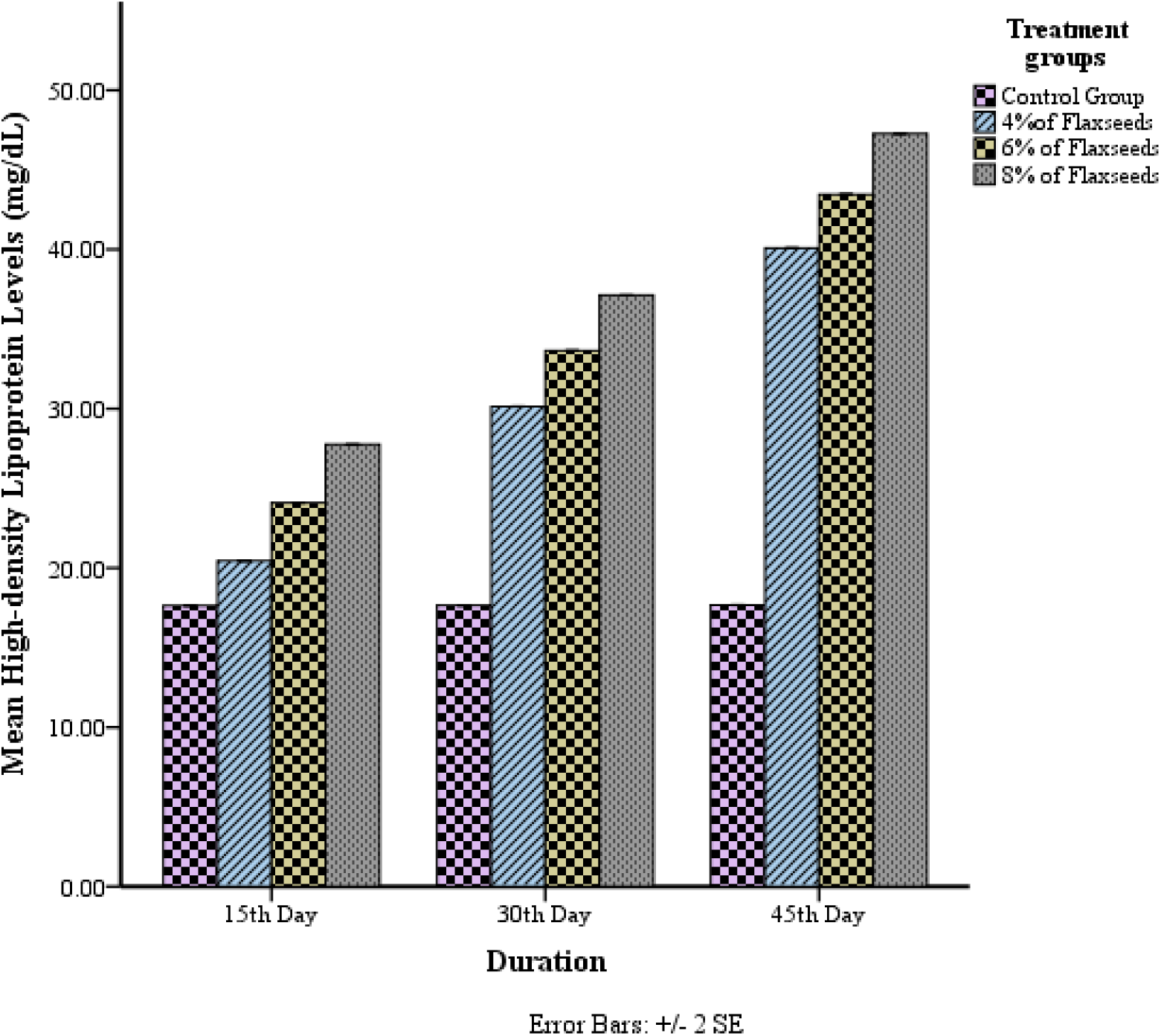
Concentration and time dependent effect of flaxseeds on high-density lipoprotein (HDL) level (mg/dL) of rabbits in T_0_ (Control), T_1_ (4% flaxseeds), T_2_ (6% flaxseeds) and T_3_ (8% flaxseeds) groups at different time-scales (15th, 30th and 45th days).

### Effect of flaxseeds on immunity

Treatment of rabbits with 4%, 6% and 8% of flaxseeds for 45 days resulted in significant (p≤0.05) rise in antibody titer in all treated groups T_1_ (4% flaxseeds), T_2_ (6% flaxseeds) and T_3_ (8% flaxseeds) in comparison to T_0_ (control group). The consumption of flaxseeds depicted maximum increase in the immunity of rabbits at concentration 8% of flaxseeds (Figure-18) with mean antibody titer value of 5.98±.00 mg/dL (Table-4).

**Figure 18.**
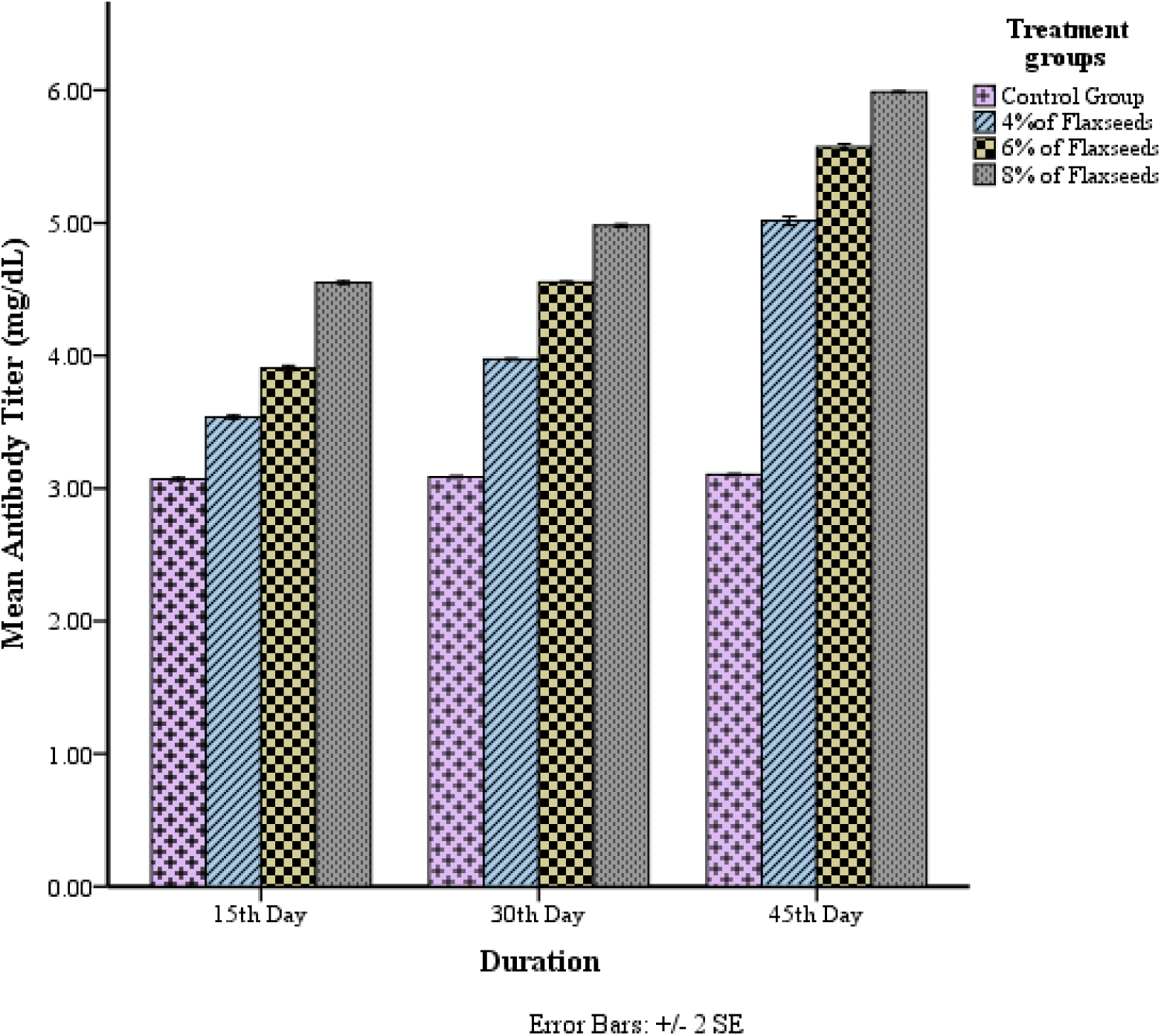
Concentration and time dependent effect of flaxseeds on antibody titer (mg/dL) against rabbit hemorrhagic disease virus (RHDV) of rabbits in T_0_ (Control), T_1_ (4% flaxseeds), T_2_ (6% flaxseeds) and T_3_ (8% flaxseeds) groups at different time-scales (15th, 30th and 45th days).

**Table 4.** Effect of Flaxseeds on Antibody Titer (mg/dL) of Rabbits at 15^th^, 30^th^ and 45^th^ Day (Mean±S.E.M)

| **Duration**<br>**(in days)** | **Treatments** |  |  |  |
| --- | --- | --- | --- | --- |
|  | **T<sub>0</sub> (Control)** | **T<sub>1</sub> (4% of**<br>**flaxseeds)** | **T<sub>2</sub> (6% of**<br>**flaxseeds)** | **T<sub>3</sub> (8% of**<br>**flaxseeds)** |
| 15<sup>th</sup> | 3.07±.00<sup>d</sup> | 3.53±.00<sup>c</sup> | 3.90±.00<sup>b</sup> | 4.55±.00<sup>a</sup> |
| 30<sup>th</sup> | 3.08±.00<sup>d</sup> | 3.97±.00<sup>c</sup> | 4.55±.00<sup>b</sup> | 4.98±.00<sup>a</sup> |
| 45<sup>th</sup> | 3.10±.00<sup>d</sup> | 5.01±.01<sup>c</sup> | 5.57±.01<sup>b</sup> | 5.98±.00<sup>a</sup> |

*Means with different superscript in a column are significantly different from one another (p≤0.05) Tuckey's Test.*

## Discussion

### Hematological parameters

The results were compatible with a study that showed significant positive effect on different hematological parameters when flaxseeds were added to the diet of rabbits. These parameters were erythrocyte indices, HCT, RBC, MCHC and MCH values [21]. Another study indicated that flaxseed powder and oil reversed the hematological abnormalities in rats suffering from T2DM [18].

Flaxseeds are rich with phytoestrogens as ligninss and secoisolariciresinol diglucoside (SDG) is the most essential phytoestrogen in flaxseeds [20]. Consumption of lignins complex (extracted from flaxseeds) reduced the impacts of hypercholesterolemic atherosclerosis by lowering the concentrations of serum lipids in the rabbits. However, in hypercholesterolemic and normocholesterolemic rabbits, the short term use of lignins complex did not cause any negative effect on the concentrations of various blood parameters like MCV, RBCs, lymphocytes, WBCs, Hb, monocytes and platelets [24].

### Lipid profile

Following results were in accordance with an investigation where cholesterol lowering effects were depicted in rabbits. The intervention of whole flaxseed in rabbits indicated reduction in TC and LDL-C levels but no changes were observed while using flaxseed oil. The reduction in the TG probably occur due to the presence of PUFA n-3 in the flaxseeds [23, 12]. Similarly, another study reported the significant improvement in the serum parameters of mice fed with high-fat diet SDG (0.5 and 1% w/w) for 4 weeks. Administration of 1% SDG to the mice’s diet showed significant fall in TG and TC [16].

Flaxseeds have positive influence on lipid profile due the presence of ALA that can significantly lower TC, LDL and VLDL cholesterol [5, 13]. Flaxseeds possess essential bioactive elements like fatty acids, proteins, lignins, vitamins, dietary fibers, minerals, and carbohydrates [4, 10, 13, 22]. These bioactive elements make flaxseeds exhibit properties such as anti-oxidant, anti-inflammatory, antibacterial etc [1, 26]. Flaxseeds have immense therapeutic effects to cope respiratory disorders, urinary tract infections, skin inflammations and treat diseases like coronary heart disease (CHD), arteriosclerosis, diabetes, and lupus nephritis [3, 6, 29].

### Immunity

An investigation conducted on 45 white New Zealand rabbits (male) divided into 2 groups diabetic animal and non-diabetic group reported the positive effect of flaxseeds on their immunity. Four wounds (linear shape full thickness) were made in each animals on both sides of their backbone skin. Results indicated significant reduction in infiltration of inflammatory cells at 14^th^ day and prominent change in the closure rate of wound in the study groups that consumed flaxseeds (diabetic as well as non-diabetic). High re-epithelization potential reported in both groups. High neovascularization rate reported in diabetic group of animals that received flaxseeds as compared to control diabetic group [2].

Similarly, eight rabbits in each group fed with flaxseeds, containing abundant amount of polyunsaturated fatty acids, flaxseeds plus hazelnut skins (with antioxidant property) or palm oil, (containing high amount of saturated fatty acids) reported increase in the blood value of lysozyme. Lysozyme value increased significantly in group taking flaxseeds in contrast to other groups. Lysozyme is an immune parameter associated to the phenomena of inflammation [8].

Polyunsaturated Fatty Acid (PUFA) present in flaxseeds positively enhances the immune profile of dairy ewes. FS (flaxseeds) given to ewes during or around parturition, can positively influence immune response by changing the cytokine production. The results indicated that the ewes with FS containing diet, their IL-6 level remained the same up to 14 days of postpartum and then decreased from 14 to 42 day of postpartum. At 14^th^ day, the ewes fed with FS showed significant rise in IL-10. During parturition, ewes fed with FS showed significant fall in IL-1β as compared to ewes in control group [9].

Anti-inflammatory and immunoregulatory role of flaxseeds is mostly due to excessive amount of ALAs or n-3 PUFAs present in them [31]. ALA plays a role in safeguarding blood vessels from inflammatory damage. The incorporation of dietary flaxseeds can elevate blood levels of ALA and thus reduces inflammation [17].

## Conclusion and recommendations

The research work depicted the significant (p≤0.05) effect of different concentrations of flaxseeds on lipid profile, hematological parameters and immunity of male rabbit. The essential bioactive components in flaxseeds induced positive changes in lipid profile, hematological parameters and immunity of rabbits in all treated groups comparable to the control group. Significant reduction in LDL-C, TC and TG and and rise in HDL-C indicated positive influence of flaxseeds on lipid parameters of rabbits. All hematological parameters also indicated significant fluctuations in all treated groups after the consumption of flaxseeds. The significant rise in antibody titer represented the significant positive effect of the seeds on the immunity of rabbits. The effect of flaxseeds was prominent in T_3_ that received high dose of flaxseeds (8%) followed by medium (6%) and low (4%) doses. Maximum values of all the parameters of lipid profile, hematological parameters and immunity were observed at concentration 8% flaxseeds.

Anti-oxidant, anti-inflammatory, immunoregulatory and anti-bacterial properties of flaxseeds can provide a number of therapeutic and nutritional benefits. The nutritive bioactive elements in flaxseeds like SDG, n-3 PUFAs, n-6 PUFAs, proteins, vitamins, fibers etc. can also make these seeds one of the best natural alternative to antibiotics. This study, therefore, suggests that the utilization of a specific concentration of flaxseeds as nutritional supplement can improve the health without any side effects.

## Data availability

All the relevant data are within the manuscript.

## Acknowledgements

Not applicable.

## Author’s contributions

Conceptualization, Methodology, Writing – original draft: Ayesha Kanwal

Project administration, Supervision, Validation: Razia Iqbal

Manuscript review, Writing – review and editing: Farah Saleem

Data curation, Formal analysis: Amna Kanwal

